# Single-cell metabarcoding reveals biotic interactions of the Arctic calcifier *Neogloboquadrina pachyderma* with the eukaryotic pelagic community

**DOI:** 10.1101/2020.10.20.347930

**Authors:** Mattia Greco, Raphaël Morard, Michal Kucera

## Abstract

Isotopic and trace-element signals in the calcite shells of the planktonic foraminifera *Neogloboquadrina pachyderma* represent key proxies to reconstruct past climatic conditions in northern high latitudes. A correct interpretation of these chemical signals requires knowledge of the habitat and trophic interactions of the species. Direct observations on the biological interactions of *N. pachyderma* in polar environments are lacking and to date no consensus exists on the trophic behaviour of this species. Here we use single-cell metabarcoding to characterise the interactions of 39 specimens of *N. pachyderma* from two sites in the Baffin Bay with the local eukaryotic pelagic community. Our results show that the eukaryotic interactome of the foraminifera is dominated by diatoms, accounting for > 50% of the reads in 17 of the samples, but other groups such as Crustacea and Syndiniales are also present. The high abundance Syndiniales suggests that these parasites could infect *N. pachyderma* and may play an important role in its population dynamics. Moreover, the strong but taxonomically non-specific association with algae, existing irrespective of depth and occurring in specimens collected far below the photic zone indicates that opportunistically grazed diatom-fuelled marine aggregates likely represent the main interaction substrate of *N. pachyderma*.

## Introduction

*Neogloboquadrina pachyderma* is the dominant planktonic foraminifera species in high latitudes where it makes up to 90% of the total assemblage (Volkmann, 2000; Schiebel *et al.*, 2017). Paleoceanographers use the geochemical signal preserved in the calcite shells of this species to investigate past states of the Arctic Ocean and reconstruct past circulation, sea ice formation and glacier meltwater events (e.g., Hillaire-Marcel et al., 2008; Knies & Vogt, 2003; Stein et al., 1994). The correct interpretation of the paleo-reconstructions relies on a thorough understanding of species-specific ecology of living planktonic foraminifera in the water column (Ezard *et al.*, 2015; Jonkers and Kucera, 2015). Most of the efforts in understanding the niche of planktonic foraminifera species have focussed on constraining the role of abiotic factors (e.g., Lessa et al., 2020; Rebotim et al., 2017), but it is increasingly clear that biotic interactions are also important in shaping pelagic protist communities (Biard and Ohman, 2020; Lima-Mendez *et al.*, 2015).

In particular, constraining the biological interactions (diet, presence of symbionts/parasites), can dramatically improve our understanding of the mechanisms and the context of how the environmental signal is recorded in foraminifera shells (Morard *et al.*, 2019; Fehrenbacher *et al.*, 2018). In addition, understanding the biotic interactions of the species can help develop more accurate numerical models that predict the response of *N. pachyderma* to future climate change (Kretschmer *et al.*, 2018; Roy *et al.*, 2015).

Despite being widely applied in palaeoceanography, the ecology of *N. pachyderma* in the Arctic remains elusive (Xiao *et al.*, 2014). A recent pan-Arctic investigation on the distribution of this species highlighted the necessity to disentangle its biological interactions as abiotic factors alone could only explain a fraction of the observed variability in its habitat depth (Greco *et al.*, 2019). Besides speculations based on cytoplasm pigmentation (Kohfeld and Fairbanks, 1996; Stangeew, 2001), no direct observations currently exist on the diet of *N. pachyderma* in the Arctic Ocean (Volkmann, 2000; Bjorbækmo *et al.*, 2020). Most authors consider this species as herbivorous (Kohfeld and Fairbanks, 1996; Manno and Pavlov, 2014; Pados and Spielhagen, 2014; Schiebel *et al.*, 2017), other as omnivorous (Stangeew, 2001), while in culturing experiments the species is known to survive when fed with live or dead *Artemia*, and therefore could be regarded as carnivorous (Manno *et al.*, 2012). Next to the food source, other biotic interactions of *N. pachyderma* have not been investigated, and nothing is known about interactions such as presence of symbionts or parasites (Bjorbækmo *et al.*, 2020). Such observations could be relevant for the understanding and prediction of a range of pelagic processes, including the carbon cycle in the Arctic Ocean. For example, by increasing physiological stress on and possibly killing *N. pachyderma*, parasite activity could affect the downward flux of dead cells which would result in the release of particulate organic matter (POM), fuelling in turn the microbial loop (Skovgaard, 2014).

Gaining insights on Arctic protistan trophic interactions has been historically challenging as it used to require well-developed culturing protocols and time-consuming microscope observations. The advent of high-throughput sequencing presents the opportunity to overcome such limits (Lovejoy, 2014; Prazeres *et al.*, 2017; Bird *et al.*, 2017, 2018). In this study, we use single-cell metabarcoding to constrain trophic interactions between *N. pachyderma* and the eukaryotic pelagic community in the Baffin Bay. We focus on the eukaryotic organisms (eukaryome) as previous work (Bjorbækmo *et al.*, 2020) and existing data on feeding behaviour in planktonic foraminifera indicate that biotic interactions are likely to be mainly with eukaryotes (Schiebel and Hemleben, 2017).

To this end, we identify the taxonomic composition of eukaryomes extracted from 39 *N. pachyderma* specimens collected at two different depths from two sites representing distinct oceanographic settings (Fig. 1). To identify the interacting pelagic community, we use the classical metabarcoding approach to sequence bulk DNA extracted from contextual seawater samples. We use the resulting dataset to test the specificity of *N. pachyderma* interactions with the eukaryotic community and their ecological significance by comparing (i) the data derived from single-cell metabarcoding and from water samples (ii) the taxonomic composition in specimens collected at different depths, and (iii) the taxonomic composition in specimens sampled at two locations with contrasting trophic conditions.

**Figure 1.**
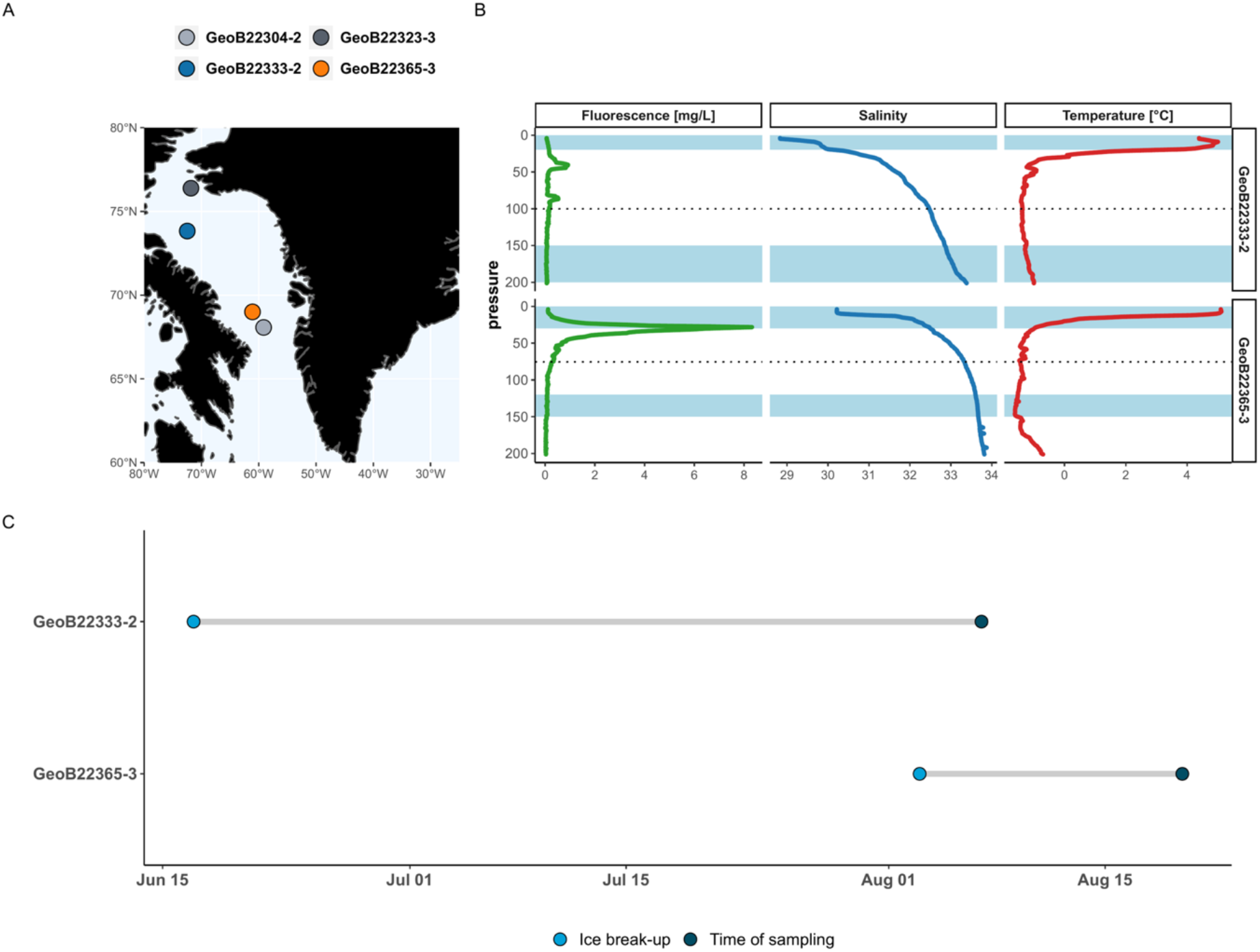
A) Sampling locations. B) CTD vertical profiles of fluorescence (productivity), salinity and temperature at the two stations where planktonic foraminifera specimens were analysed. Blue shaded areas represent the sampled depth intervals and the dotted line the limit of the Euphotic zone (calculated using the approach described in Lee *et al.*, 2007). C) Time between ice break-up and sampling date at the two studied sites.

## Material and Methods

### Sampling

In August 2017, during the MSM66 cruise in the Baffin Bay on the R/V Maria S. Merian, planktonic foraminifera were sampled at different depths using a Multi-net (HydroBios 92B, Kiel, Germany), equipped with five net bags with 100 μm mesh diameter. Individual specimens were isolated from the plankton sample and stored on micropaleontological slides at a temperature of −80°C. In parallel, seawater (1L) from Niskin bottles was sampled at surface (~5 m) and at depth (200 m) and filtered through 0.2 μm cellulose filters. Filters were then stored in buffer (1.8 mL of 50mM Tris–HCl,0.75M sucrose and 40mM EDTA; pH 8.3) at a temperature of −80°C. Water samples were collected from two sites in the North Baffin Bay (GeoB22323-3 and GeoB22333-2) and two sites from the south of the Baffin Bay (GeoB22365-3 and GeoB22304-2) (Fig.1). *N. pachyderma* specimens collected from different depth at one of the sites in the north, GeoB22333-2 (sampling intervals: 0-20 m; 150-200 m), and from one in the south, GeoB22365-3 (sampling intervals: 0-30 m; 120-150 m) (Fig. 1), were selected for single-cell metabarcoding analyses.

### Environmental data

At each sampling station, a conductivity– temperature–depth (CTD) device equipped with a fluorescence sensor (WETLabs ECOFLNTU (RT)D) was deployed to obtain vertical profiles of physical properties and algae pigment concentrations (Dorschel *et al.*, 2017). The time of sea-ice break-up at the two stations was determined by extracting in situ sea-ice concentration from 25km×25km resolution passive microwave satellite raster imagery obtained from the Sea Ice Index Version 3.0 product of the National Snow and Ice Data Centre using a custom script in R (Fetterer *et al.*, 2017; R Core Team, 2017).

### DNA Extraction, Amplification and Sequencing

Specimens of *N. pachyderma* were transferred from the slides and total DNA was extracted from each specimen following the GITC* protocol (Weiner *et al.*, 2016). Total DNA from filters was extracted using E.Z.N.A. kit following manufacture instructions including blank extractions to control for (cross-) contamination events. DNA extractions were then amplified in triplicates using the universal Eukaryotic V4 tagged primers TAReuk454FWD1 (5’-CCAGCA(G⁄C)C(C⁄T)GCGG-TAATTCC-3’) and TAReukREV3 (5’-ACTTTCGTTCTTGAT(C⁄T)(A⁄G)A-3’) (Stoeck *et al.*, 2010) that amplify only eukaryotes and offer a good taxonomic resolution across the entire eukaryote realm (del Campo *et al.*, 2019). Each tagged PCR primer consists of a unique tag sequence of 8 nucleotide appended to the 5’end of the common amplification primer sequence.

Each PCR was performed in a total volume of 25 μl, including 0.02 U/ μl of Taq DNA polymerase (Phusion), 1.03 umol/μl of 5 × Green Buffer (Phusion), 0.2 mM of each dNTP, 0.41 umol/μl of each primer, 2.56 umol/μl of MgCl_2_ and 1 μl of DNA extract. The conditions for the first amplification consisted of a pre-denaturation step at 98 °C for 30 s to melt the complex genomic DNA mixture, followed by 35 cycles of denaturation at 98 °C for 10 s, annealing at 52 °C for 30 s and extension at 72 °C for 45 s, followed by a final extension step at 72 °C for 10 min.

Positive triplicates were successively purified using the QIAquick PCR purification Kit (Qiagen) in a final volume of 30 μl, and the DNA content quantitated using a QUANTUS fluorometer (Promega). Samples were finally loaded in 4 different pools with each unique tagged primer combination present only once in each pool. The purified products from the water samples were pooled with twice the amount of DNA than the single-cell foraminifera to get a higher sequencing depth and obtain a better representation of the diversity in the environment. Pools were sent to the University of Geneva for sequencing Illumina (MiSeq) where sequencing libraries were prepared using the reagents of the PCR-free TruSeq kit (Illumina).

### Sequence data pre-processing and taxonomic assignment

In total, amplicon sequencing produced 10,793,476 raw paired-end (PE) reads. Raw reads were demultiplexed with Cutadapt 2.7 (Martin, 2013) using the Combinatorial Dual Indexing option allowing for 2 bp errors in the barcode sequence and no indels. Reads with the valid barcode combinations were selected for the following steps and reads containing ambiguous bases (Ns) were removed using DADA2 1.14.1 (Callahan *et al.*, 2016) in R 3.6.0. Primers were removed from the reads using the “linked” adapter option in Cutadapt 2.7. At most 2 errors were allowed during filtering, while 20 bp were trimmed from the end of the forward and 50 bp from the end of the reverse reads to remove barcodes and primers. Reads were further processed with the DADA2 pipeline in R 3.6.0. The minimum allowed read length was set to 175 bp. After dereplicating forward and reverse reads, the DADA2 pipeline was used to identify amplicon sequence variants (ASVs) in the dataset. The forward and reverse reads were merged and chimeras were identified and removed based on matches with combinations of 3’- and 5’-segments of different sequences. The ASVs were then taxonomically classified with the naïve Bayesian classifier method implemented in DADA2 based on the PR^2^ database, a curated reference 18S rRNA database spanning the eukaryotic tree of life (Guillou *et al.*, 2013; del Campo *et al.*, 2018).

### Statistical Analyses

After filtering steps, our dataset was reduced to 4,035,059 reads. Because we focus on the direct eukaryotic interactions in our analyses, 88 ASVs belonging to land plants and land mammals were considered as contaminants and removed from our dataset.

For downstream statistical analyses, sample triplicates were merged and the dataset was then rarefied to the minimum sampling depth (24,652 sequences per sample) (Fig. S1) using the phyloseq package in R (McMurdie and Holmes, 2013). The differences in the taxa composition in the foraminifera samples were explored using a multivariate approach. The ASVs abundance were binned at genus level and transformed in relative proportions. From the resulting dataset, we calculated Bray Curtis dissimilarity among the different samples and visualised the similarity structure by two-dimensional Principal Coordinates Analysis (PCoA). Next, in order to test whether collection depth or sampling site significantly affected the interactome composition of individual foraminifera, we performed a permutation multivariate analysis of variance (PERMANOVA) using the adonis function in the vegan R package (Oksanen *et al.*, 2018). Finally, to understand the differences in the composition between groups, we carried out the Analysis of Composition of Microbiomes (ANCOM) using the ANCOM package in R (Mandal *et al.*, 2015; Kaul *et al.*, 2017). This test performs a differential abundance analysis on the ASV table (not binned and not transformed) to detect differentially abundant ASVs across different experimental group allowing to control for False Discovery Rate (FDR); we opted for a high cut-off (0.9) for a conservative interpretation of the results (smaller FDR). When no significant difference between the experimental groups was observed, the test was repeated with a lower cut-off (0.7).

## Results

A total of 1890 ASVs were observed in the dataset. The amount was reduced to 340 ASVs when reads were collapsed at genus level. Crustaceans (Class Arthropoda) and Dinoflagellates (Class Dinophyceae) dominated the community in the water samples across different sites and depth.

The fact that the sampling sites GeoB22365-3 and GeoB22333 presented quite similar thermohaline profiles but differed in terms of productivity (Fig.1). This was reflected in the taxonomic signal of the water samples showing a higher proportion of ASVs belonging to diatoms (Class Bacillariophyta) at the GeoB22365-3 station, especially in the surface layer (Fig. 2).

**Figure 2.**
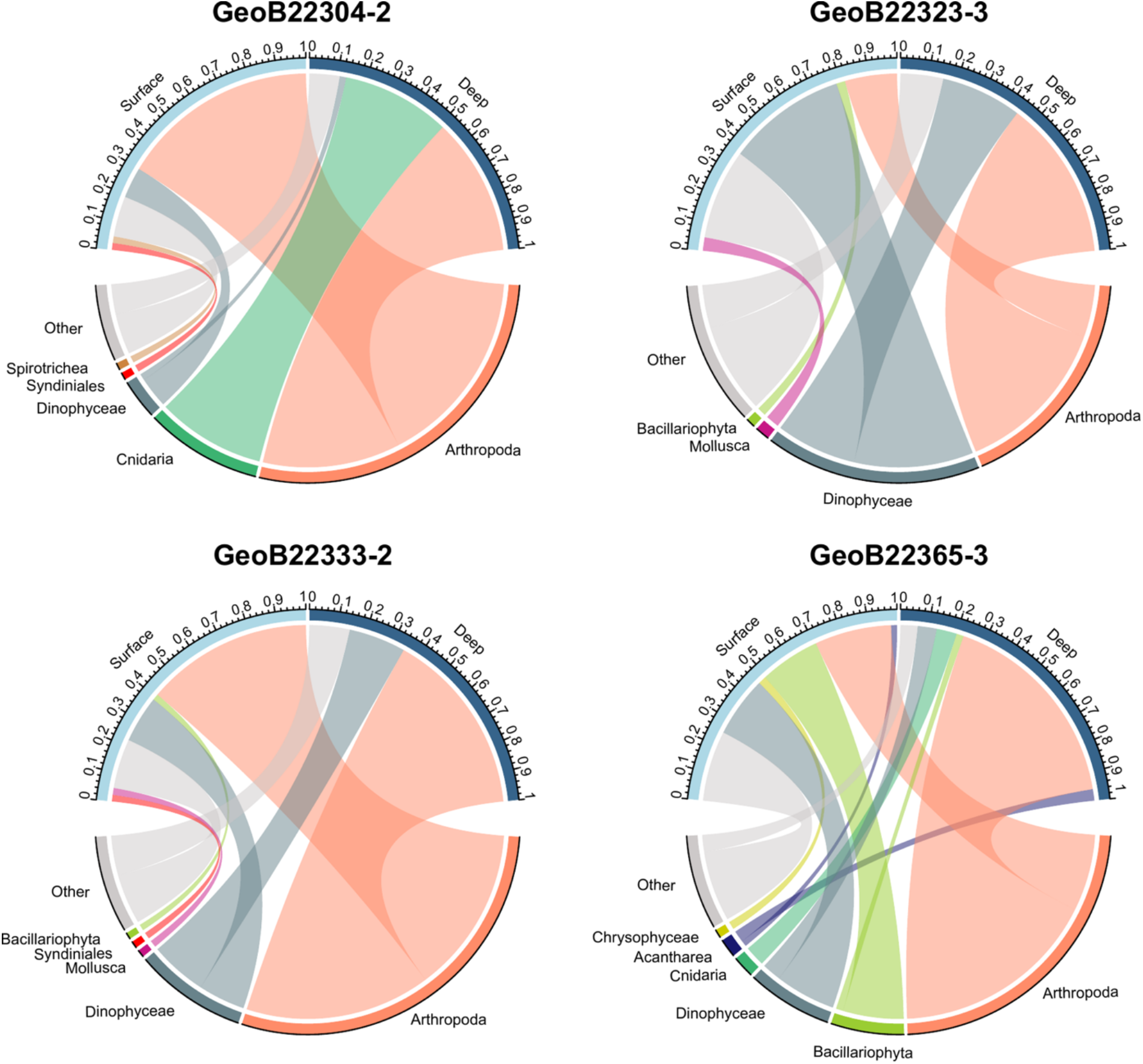
Taxonomic composition of the amplicons obtained from DNA extracts retrieved from filtered ambient water samples at the surface and subsurface. Colours represent different taxonomic groups (taxonomic groups occurring with a frequency below 0.01 were condensed in the category “Minor Taxa”).

In the foraminifera samples, diatom ASVs (Class Bacillariophyta) dominated the eukaryome in specimens from the surface layer (Fig.3), reaching abundance above 50% in 17 specimens. The tree maps in Fig. 4 show that within the Bacillariophyta, the genus *Chaetoceros* was the most abundant in both water and foraminifera samples especially from site GeoB22333, represented by ASVs belonging to the cold-water species *Chaetoceros gelidus* and *Chaetoceros socialis*. In the *N. pachyderma* specimens from the GeoB22365 site, a higher proportion (30%) of the raphid-pennate diatoms of the genus *Fragilariopsis* was present. The algal composition in the water sample from the same site was more diverse with abundant ASVs from the genera *Pseudo-nitzchia* and *Thalassiosira*. Furthermore, the foraminifera collected at station GeoB22365 yielded a high number of ASVs belonging to Syndiniales representing up to 40% of the total assemblage in the specimens sampled at deeper depth (Fig. 3).

**Figure 3.**
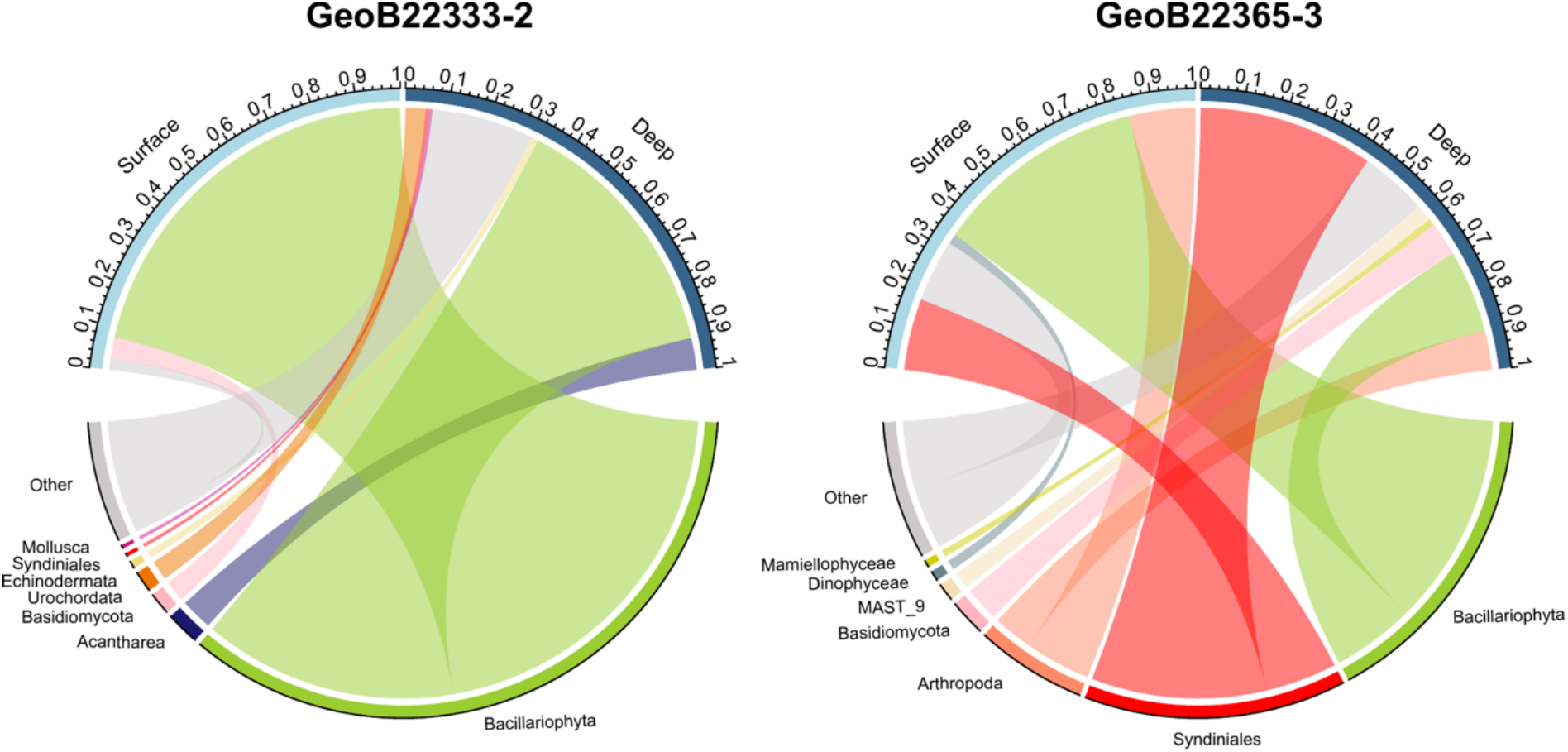
Taxonomic composition of the amplicons retrieved from single-cell extractions from *N. pachyderma* sampled at different depths. Colours represent different taxonomic groups (Taxonomic groups occurring with a frequency below 0.01 were condensed in the category “Minor Taxa”).

**Figure 4.**
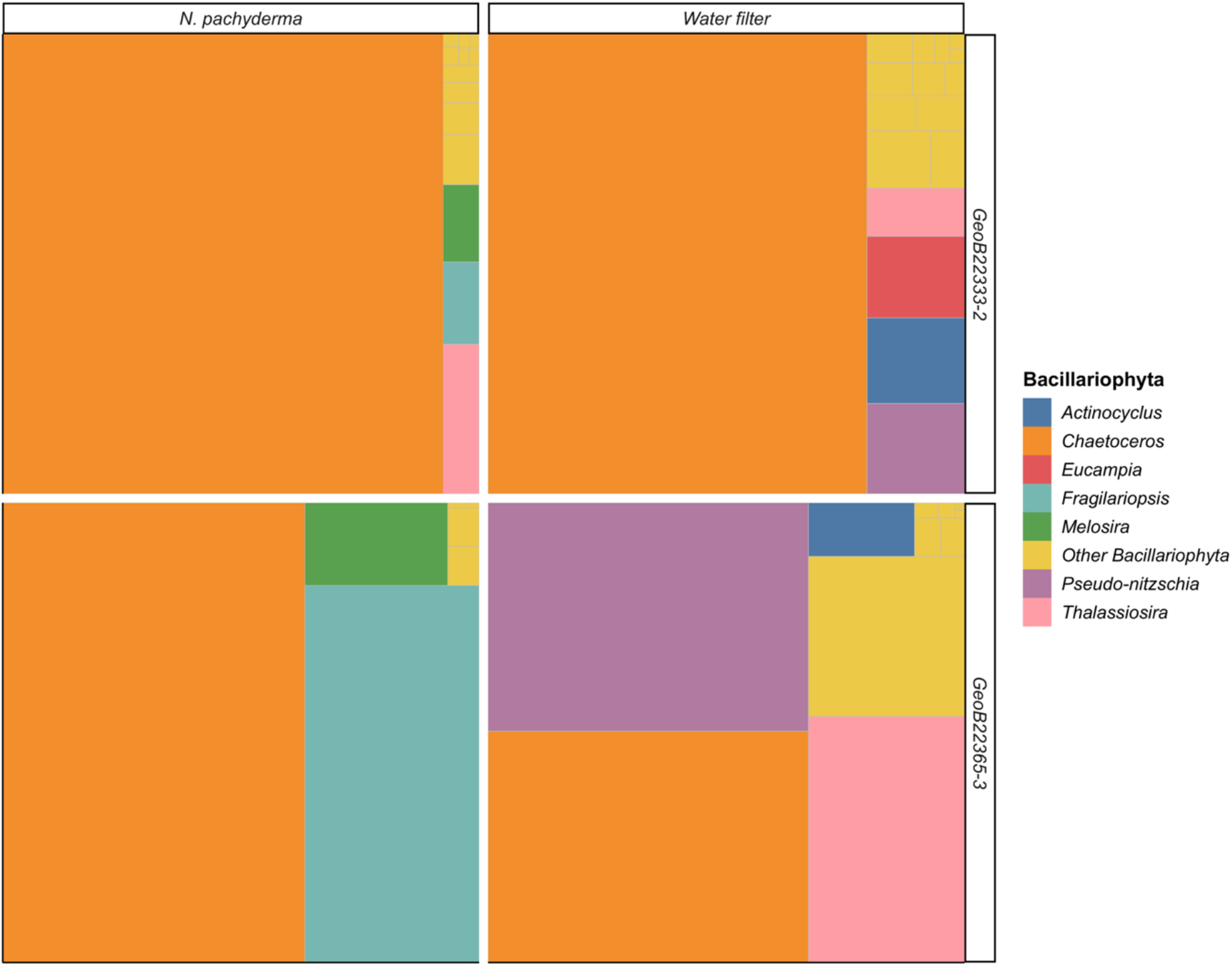
Treemaps showing the average Bacillariophyta amplicon composition in foraminifera and in the ambient water samples averaged throughout the water column at the two sites. Colours represent the different identified genera (Taxonomic groups occurring with a frequency below 0.01 were condensed in the category “Other Bacillariophyta”).

The Principal Coordinate Analysis (PCoA) of the foraminifera eukaryomes indicated systematic differences in composition by site and also by depth (Fig. 5). The majority of the variance (50.8 %) was explained by the first component separating eukaryomes from the two stations irrespective of sampling depth. The PERMANOVA analyses confirmed that the differences in community linked to sampling site was statistically significant (*p-value* = 0.001, R^2^= 32%). A much lower, but still significant (*p-value* < 0.05, R^2^= 6%), portion of variability in the composition was explained when collection depth was tested as grouping factor (Table 1). Individual specimens collected at the same depth differed in the composition of their eukaryome, with higher variability among specimens from deep layer at the station GeoB22333-2 and among specimens from the surface layer at station GeoB22365-3. (Fig. 5).

**Table I.** PERMANOVA results based on Bray-Curtis dissimilarities using genus abundance in the foraminifera samples in relation to compartment for a) Sampling site and b) depth of collection (*p-values* based on 999 permutations).

| | Df | Sum Sq | Mean Sq | F model | $R^2$ | *p-value* |
| --- | --- | --- | --- | --- | --- | --- |
| **a) <u>Station</u>** | 1 | 3.221286 | 3.221286 | 17.42954 | 0.320222 | 0.001 |
| Residuals | 37 | 6.838252 | 0.184818 |  | 0.679778 |  |
| Total | 38 | 10.05954 |  |  | 1 |  |
| **b) <u>Depth</u>** | 1 | 0.65178 | 0.65178 | 2.563399 | 0.064792 | 0.044 |
| Residuals | 37 | 9.407759 | 0.254264 |  | 0.935208 |  |
| Total | 38 | 10.05954 |  |  | 1 |  |

**Figure 5.**
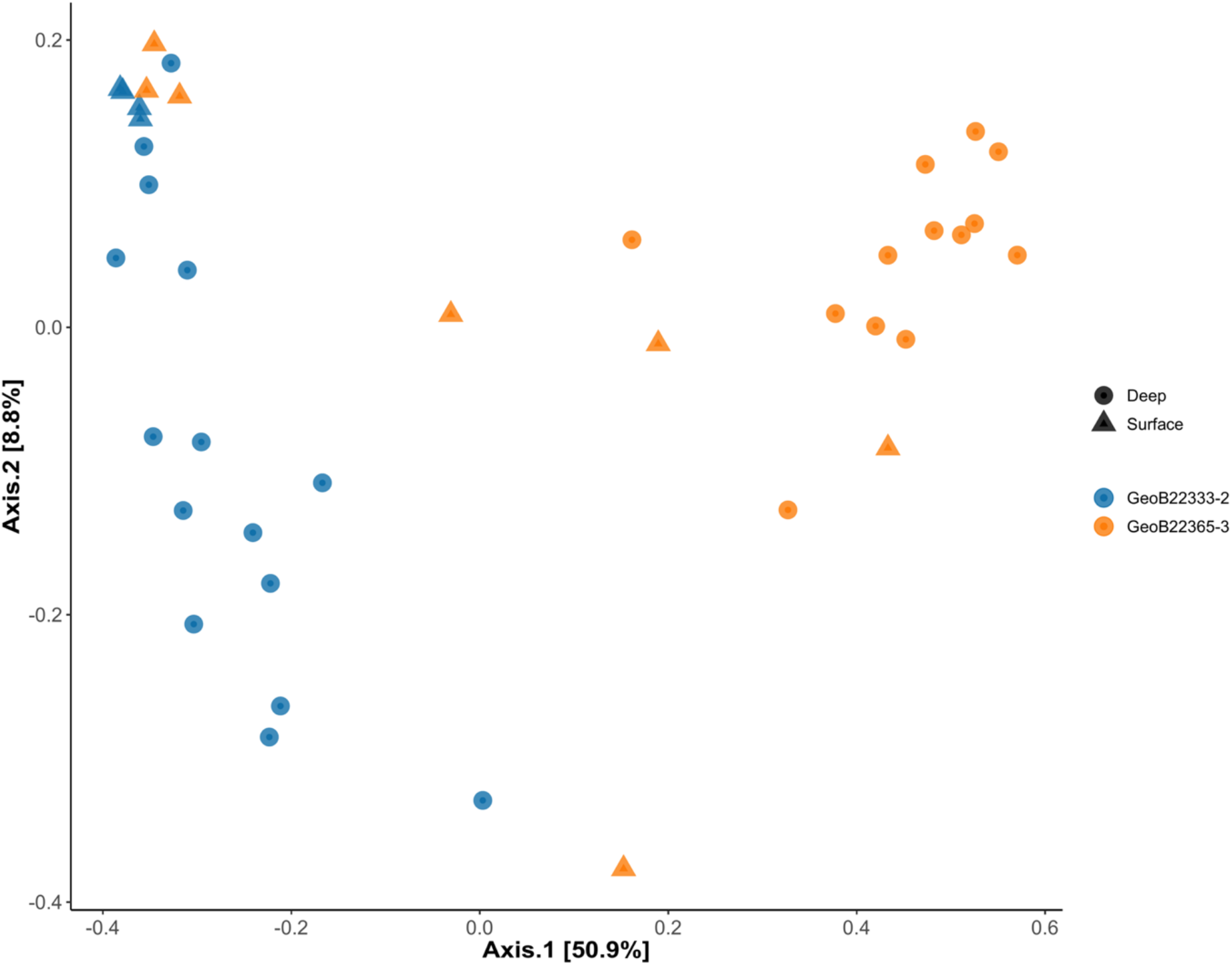
PCoA of Bray-Curtis dissimilaritiy score calculated on genus relative frequency of all taxa present in each *N. pachyderma* sample. Colours represent sampling site and shapes the depth of collection.

The ANCOM analysis indicated that ASVs belonging to diatoms (*Chaetoceros*), Syndiniales (*Dino Group I*) and Acantharians (*Chaunacanthida*) were significantly different in abundance between the two sampling sites (Fig. 6). On the other hand, no significant difference in ASV abundance was found between specimens collected at different depth, when the same FDR threshold was used. Relaxing the threshold to 0.7 resulted in the identification of two ASVs of the genus *Chaetoceros* as being significantly more abundant in the shallower *N. pachyderma* specimens.

**Figure 6.**
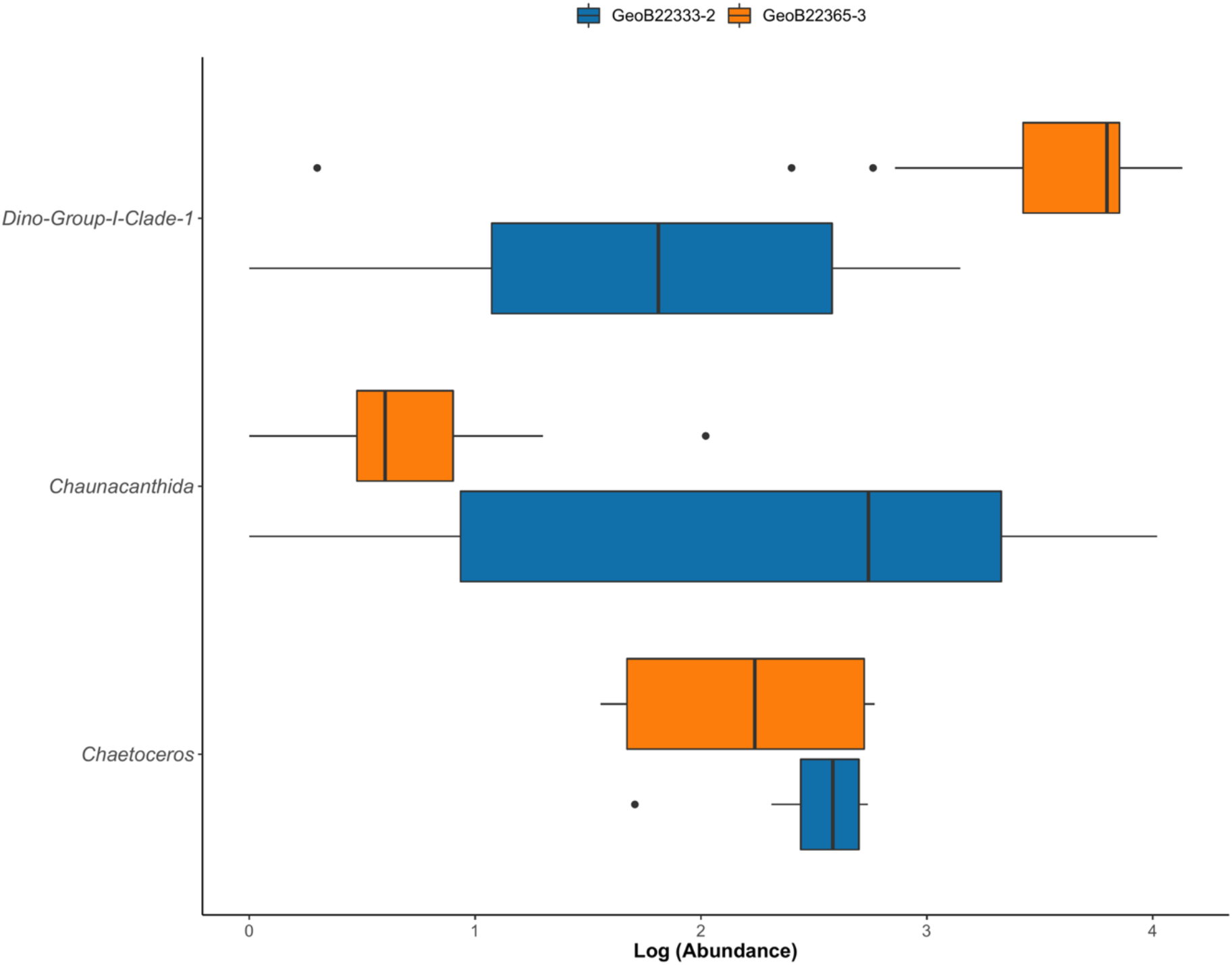
ASVs found in the ANCOM analysis (abundance is log transformed) to be associated with differences between the two sampling sites.

## Discussion

The pelagic community signal recovered from the water samples appeared homogenous in terms of the main taxonomic groups represented (Fig. 2) but a deeper look at the algal composition reveals a signature of the different ecological conditions at the two sites at the time of collection (Fig. 4). At station GeoB22333, most of the Bacillariophyta ASVs in both water and foraminifera samples were assigned to the two closely related species *C. gelidus* and *C. socialis*, known to be the most abundant centric diatoms in the Baffin Bay during summer (Chamnansinp *et al.*, 2013; De Luca *et al.*, 2019). In the water samples from site GeoB22365, along with *Chaetoceros*, another centric diatom, *Thalassiosira* is highly represented. Blooms of diatoms of the genus *Thalassiosira* have been described as intense and transient, and are usually rapidly replaced by *Chaetoceros* spp. blooms (Booth *et al.*, 2002; Lafond *et al.*, 2019).

Next to *Chaetoceros* and *Thalassiosira*, diatoms of the genus *Pseudo-nitzchia* are the most abundant in water samples from station GeoB22365. This pennate diatom taxon is generally observed in locations where sea ice cover is present (Poulin *et al.*, 2011) and usually precedes *Thalassiosira* and *Chaetoceros* in the algal bloom succession after sea-ice break up (Lafond *et al.*, 2019). In *N. pachyderma* specimens collected at the same station, the most abundant Bacillariophyta ASVs belonged to sea-ice diatom *Fragilariopsis* (Mock *et al.*, 2017) and *Melosira arctica,* a diatom that grows filaments anchored on the underside of the sea-ice (Poulin *et al.*, 2014; Boetius *et al.*, 2013).

The algal taxonomic composition of both water and foraminifera datasets combined with CTD and sea-ice data (Fig. 1) indicate that sampling at the two sites occurred at different stages of the local algal bloom succession. Site GeoB22365 was sampled shortly (17 days) after sea-ice break with a mixed algal structure transitioning from sea-ice associated taxa to centric diatoms dominated pelagic community. The situation for the sampling at site GeoB22333 was different as here the sea ice melted 51 days before the sampling date (Fig.1c) and the algal community displayed a more homogenous structure dominated by diatoms of the genus *Chaetoceros* that are able to maintain their populations at low nutrient levels (Booth *et al.*, 2002).

The differences in the algal community structure between the two sites significantly affected the taxonomical structure of the eukaryomes in the foraminifera samples as confirmed by the PCoA and PERMANOVA results (Fig. 5, Table 1). This was expected as diatom ASVs were extremely abundant in most of *N. pachyderma* specimens. The ANCOM outcome is also consistent with a change in the algal community since the ASVs from the genus *Chaetoceros* were recognised as significantly more abundant in the specimens collected at station GeoB22333. The weaker depth-related signal of relative genus abundance detected in the PERMANOVA in the foraminifera samples, is also likely related to the diatoms with ASVs of the genus *Chaetoceros* showing a reduced abundance in deeper waters as revealed in the ‘relaxed’ ANCOM test.

Contrary to benthic foraminifera, diatoms endosymbionts have not been observed in planktonic species (Hemleben *et al.*, 1989). Moreover, a recent survey on photosymbiosis in planktonic foraminifera, have shown that the species *N. pachyderma* possesses chlorophyll but this chlorophyll is not associated with active photosynthesis (Takagi *et al.*, 2019). Given the low specificity of the interaction between *N. pachyderma* and the diatoms, we conclude that the algal ASVs recovered in the eukaryotic interactome of *N. pachyderma* likely represent the main food source of this species. Since planktonic foraminifera do not participate in daily vertical migration (Manno and Pavlov, 2014; Meilland *et al.*, 2019), the observation of similar diatom interactome compositions among specimens from the surface layer and those collected at depth far below the photic zone (Fig. 5, Table 1) allows us to conclude that at least some of the foraminifera must have been feeding on dead cells, sinking through the water column.

Diatoms of the genus *Chaetoceros* generally from chain-like structures which in the presence of high levels of biomass, tend to cluster together into larger colonies forming aggregates and producing abundant exopolymeric gels that lead to high carbon export (Chamnansinp *et al.*, 2013; Booth *et al.*, 2002; Duret *et al.*, 2020). This is particularly true in the Baffin Bay, where it has been estimated that in July, cells of *C. gelidus* can contribute up to 91 and 49% of total phytoplankton abundance and carbon respectively (Booth *et al.*, 2002). We speculate that these diatom-fuelled aggregates can represent the principal microhabitat of *N. pachyderma*.

This is consistent with the hypothesis by Fehrenbacher et al. (2018), who deduced from shell composition data that non-spinose planktonic foraminifera species like *N. pachyderma* may calcify within organic aggregates. The inference that aggregates represent the main interaction substrate of *N. pachyderma* with the pelagic community can also explain why sea-ice associated diatoms were detected in the interactome of the foraminifera. At the time of the sampling, their bloom was over in the water column, but they were represented in aggregates, which were grazed on by *N. pachyderma*, thus revealing the bloom structure as it occurred in the water column some weeks earlier.

As shown by the case of Bacillariophyta DNA in *N. pachyderma*, it is important to consider that the identification of various ASVs in the interactome does not necessarily indicate that the interaction exists when organisms are alive. With this in mind, we can interpret the non-diatom ASVs in the foraminifera samples (Fig. 3) in light of the aggregate microhabitat hypothesis. Crustaceans and soft-bodied Urochordata are also known to actively or passively participate in the formation of marine aggregates (Duret *et al.*, 2020). By feeding indiscriminately on organic particles present in the aggregates, *N. pachyderma* diet could also contain these organisms, or their remains. The ability to feed opportunistically on various components of marine aggregates would explain the diverse feeding habits of this species reported in the literature, including evidence from culturing experiments suggesting that species of this genus can digest crustaceans (Manno *et al.*, 2012; Hemleben *et al.*, 1989) as well as algae (Greco *et al.*, 2020). The higher abundance of Acantharians ASVs at site GeoB22333 is also likely associated with the sinking diatom fuelled aggregates as *Chaetoceros* is also distinctively abundant in the same site (Fig. 6). Research on Acantharians has shown that *Chaunacanthida* cysts participate in organic carbon export to depth (Decelle *et al.*, 2013), potentially explaining why their ASVs are more abundant in the deeper specimens of *N. pachyderma* (Fig. 3).

Next to diatoms, Syndiniales (group I) also constituted a large portion of the reads in *N. pachyderma* samples (Fig. 3), especially in the ones collected at Station GeoB22365 as confirmed by the ANCOM results (Fig. 6). In our dataset we could observe 10 Syndiniales (group I) ASVs in total, and for all of them the taxonomy was resolved to the Genus level.

Syndiniales are a monophyletic lineage at the base of the dinoflagellate clade, widely distributed in the world oceans (Guillou *et al.*, 2008; de Vargas *et al.*, 2015). In recent marine 18S surveys, the group I has been observed to occur in high abundance in polar oceans, in particular near the sea-ice edge and in correspondence with algal blooms (Clarke *et al.*, 2019; Cleary and Durbin, 2016; Bachy *et al.*, 2011) as we observed at sampling site GeoB22365 (Fig.1). All the Syndiniales of Group I are parasitoids that can infect distantly related hosts like other protists (dinoflagellates, cercozoans, radiolarians) or metazoans (copepods, fish eggs) and release free-living dinospores following host death (Guillou *et al.*, 2008; Clarke *et al.*, 2019). Given that Syndiniales of this group have been frequently observed in other Rhizaria (Bjorbækmo *et al.*, 2020), it is likely that *N. pachyderma* could also represent a possible host of these ubiquitous parasites. This would imply that parasitism could play a role in regulating the population dynamics of *N. pachyderma* in the Arctic. The presence of dinoflagellates parasites of the orders gymnodiniales and peridiniales in planktonic foraminifera have been previously inferred from laboratory observations (Schiebel and Hemleben, 2017) but to our knowledge, this is the first time that a parasite interaction with Syndiniales for a polar planktonic foraminifera species is inferred. However, because we cannot exclude the possibility that the Syndiniales Group I ASVs, like the diatom ASVs, represented dead specimens, our analysis alone cannot definitively resolve the putative parasite–host association. Clearly, more investigations employing laboratory observations and visualisation (e.g., the development of fluorescent in situ hybridisation probes) are needed to confirm the nature of the interaction between foraminifera and the Group I Syndiniales (Santoferrara *et al.*, 2020).

Indeed, we stress that analyses presented in this study must be seen as the first step towards the understanding of foraminifera interactome. Next to the fact that interactome based on eDNA cannot distinguish whether the interacting partners were alive or dead when present in association with the foraminifera, we also note that as long as the analysis involves PCR and only focuses on one gene, the true proportions of the interacting organisms are likely not represented correctly (Piñol *et al.*, 2015; Elbrecht and Leese, 2015) and the activity of the interacting partners cannot be resolved. However, knowing the taxonomic identity of the main partners should facilitate in the future a metatranscriptomic approach to reveal the activity of the living component of the interactome.

On the other hand, revealing consistent patterns in specimens collected at different sites and depths, in all cases distinct from the bulk seawater community, indicates that the interactome of single cells can be reconstructed from specimens collected by plankton nets and the dominance of reads likely derived from dead algal cells indicates that the DNA of foraminifera prey is preserved in their vacuoles for quite some time. However, the non-specific opportunistic feeding on aggregates that we invoke implies that an analysis carried out at one time during the seasonal cycle is unlikely to capture the full range of trophic interactions of the foraminifera and we specifically note that since our data is based on the analysis specimens larger than 100 μm, the interactions and lifestyle of *N. pachyderma* juveniles remain undetermined. Equally undetermined remains the composition and function of the prokaryotic interactome of *N. pachyderma.* Marine aggregates are prime substrate for prokaryotes (Mestre *et al.*, 2018) and the interaction of the aggregate-bound prokaryotic community with the foraminifera and/or the presence of intracellular interactions (like Bird *et al.*, 2018) would be the logical target for subsequent analyses.

## Conclusions

In this study, we used single-cell metabarcoding to constrain biological interactions between the planktonic foraminifera *N. pachyderma* and the eukaryotic pelagic community in the Baffin Bay. Diatoms (Bacillariophyta) were highly represented in most of the foraminifera samples with differences in composition that reflected the algal assemblage in the water column at the sampling site. Weaker but also significant difference in taxonomic composition of the eukaryome was observed among *N. pachyderma* specimens sampled between surface and subsurface layers. The observed non-specific relationship with pelagic diatoms, retained in specimens collected far below the photic zone, along with the presence of DNA from groups as Crustaceans and Urochordata in the foraminifera, suggest that *N. pachyderma* lives and opportunistically feeds in association with organic aggregates. In addition, our data indicate that *N. pachyderma* could be infected by Syndiniales parasites of Group I. These results advance our knowledge on the ecology of *N. pachyderma* placing it in the context of the multilevel trophic system of the Arctic pelagic realm and can improve interpretations of paleoclimatic signals preserved in its shells. Our findings showcase the potential of single-cell metabarcoding as a tool to provide important insights into planktonic microbial ecology.

## Acknowledgements

The master and crew of the F.S. Maria S. Merian are gratefully acknowledged for support of the work during the MSM66 cruise.

## Funding

This research has been supported by the Deutsche Forschungsgemeinschaft (DFG) through the International Research Training Group “Processes and impacts of climate change in the North Atlantic Ocean and the Canadian Arctic” (IRTG 1904 ArcTrain).

## Data Archiving

Raw reads will be deposited in the ENA upon acceptance. Environmental CTD data will be made available in PANGAEA (https://pangaea.de/) upon acceptance.

---

**Figure S1.**
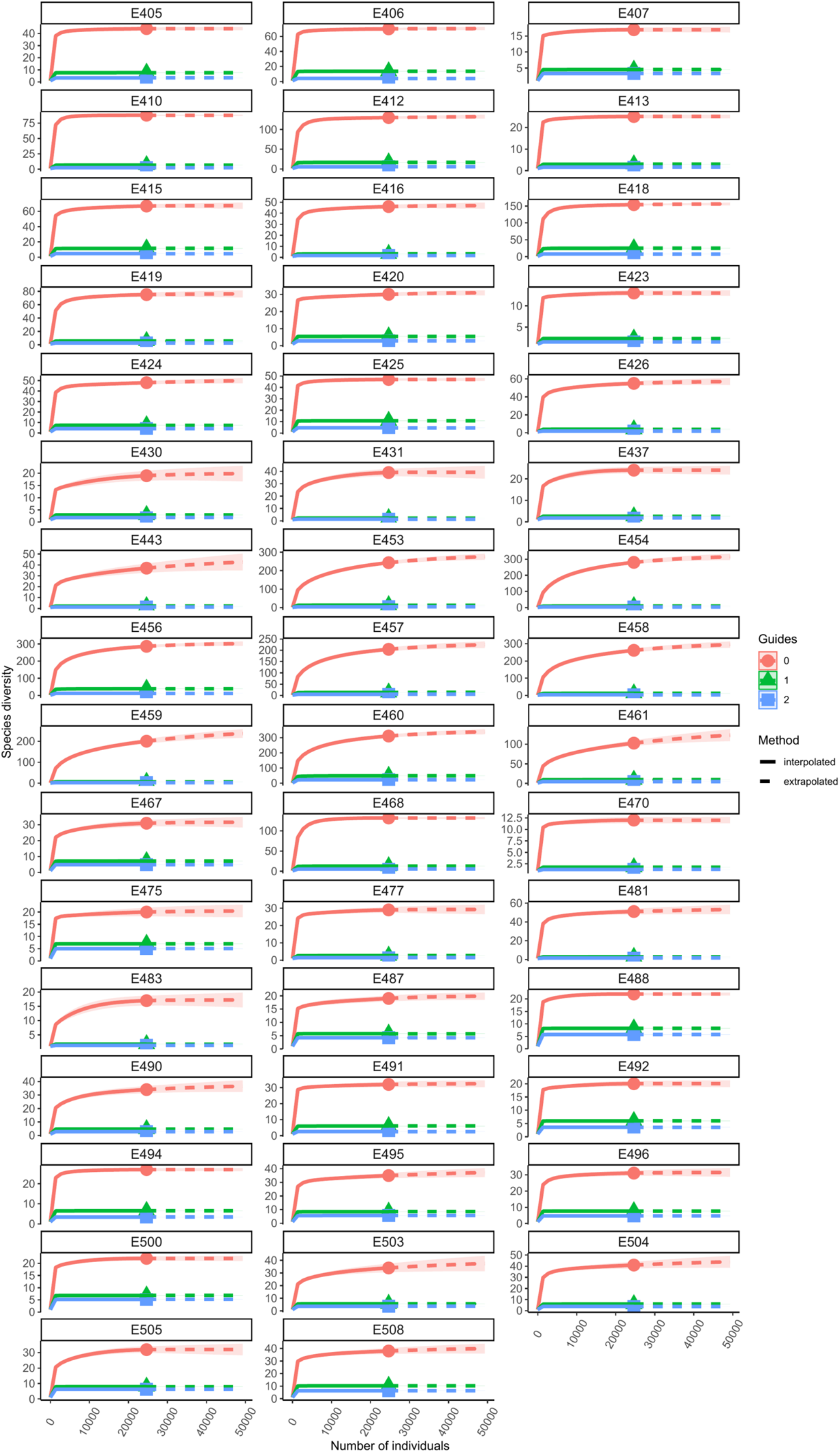
Rarefaction curves calculated using the iNEXT package in R (Hsieh *et al.*, 2016) showing (0) species richness, (1) Shannon diversity, and (3) Simpson diversity.

## Notes

### Competing Interest Statement

The authors have declared no competing interest.

